# Concurrent threats and extinction risk in a long-lived, highly fecund vertebrate with parental care

**DOI:** 10.1101/2023.12.08.570800

**Authors:** George C. Brooks, William A. Hopkins, Holly K. Kindsvater

## Abstract

Detecting declines and quantifying extinction risk of long-lived, highly fecund vertebrates including fishes, reptiles, and amphibians, can be challenging. In addition to the false notion that large clutches always buffer against population declines, the imperiled status of long-lived species can often be masked by extinction debt, wherein adults persist on the landscape for several years after populations cease to be viable. Here we develop a demographic model for the Eastern Hellbender (*Cryptobranchus alleganiensis*), an imperiled aquatic salamander with paternal care. We examined the individual and interactive effects of three of the leading threats hypothesized to contribute to the species’ demise: habitat loss due to siltation, high rates of nest failure, and excess adult mortality caused by fishing and harvest. We parameterized the model using data on their life history and reproductive ecology to model the fates of individual nests and address multiple sources of density-dependent mortality under both deterministic and stochastic environmental conditions. Our model suggests that high rates of nest failure observed in the field are sufficient to drive hellbender populations towards a geriatric age distribution and eventually to localized extinction, but that this process takes decades. Moreover, the combination of limited nest site availability due to siltation, nest failure, and stochastic adult mortality can interact to increase the likelihood and pace of extinction, which was particularly evident under stochastic scenarios. Density-dependence in larval survival and recruitment can severely hamper a population’s ability to recover from declines. Our model helps to identify tipping points beyond which extinction becomes certain and management interventions become necessary. Our approach can be generalized to understand the interactive effects of various threats to the extinction risk of other long-lived vertebrates. As we face unprecedented rates of environmental change, holistic approaches incorporating multiple concurrent threats and their impacts on different aspects of life history will be necessary to proactively conserve long-lived species.

## INTRODUCTION

Detecting declines of long-lived, highly fecund vertebrates including some fishes, reptiles, and amphibians, can be challenging (McRae et al. 2017; Cox et al. 2022). Long-lived species with high fecundity exhibit a mixture of fast and slow life history traits, thereby preventing a boilerplate evaluation of extinction risk (Hutchings et al. 2012). The imperiled status of long-lived species can often be masked by extinction debt, wherein adults persist in habitats that no longer support viable populations, and environmental perturbations or demographic shifts that undermine population viability may often go unnoticed for decades (Tilman et al. 1994, Kuussaari et al. 2009). Large clutch sizes can in theory provide resilience to extinction for such taxa, but low survival of larval stages and density-dependent recruitment often represents a key bottleneck that limits a species’ recovery potential (Wilbur 1980; Pechmann & Wilbur 1994; Alford & Richards 1999; Reynolds et al. 2005). In addition, species with complex life histories (such as distinct larval and adult stages, explosive breeding, or parental care) will often experience density dependence at multiple life stages, further reducing the capacity of populations to rebound from declines (Crouse et al. 1987; Congdon et al. 1994, Biek et al. 2002; Vonesh & De la Cruz 2002, Schmidt et al. 2005). Density-dependent processes will persist even in small or declining populations if the availability of nest sites or larval refugia have been reduced following habitat alteration. If such nuances are not incorporated into the demographic models traditionally used to evaluate population viability, the vulnerability of species to local or global extinction can be underestimated (Congdon et al. 1994, Anderson et al. 2017; Ono et al. 2019). As a result, long-lived, highly fecund vertebrates may escape the attention of conservation scientists until populations have already surpassed a gtipping point (d’Eon-Eggertson et al. 2015, de Silva et al. 2019).

Conservation efforts are further complicated by the multiple ongoing threats to biodiversity (Isaac and Cowlishaw 2004, Greenville et al. 2021, Kimmel et al. 2022). Traditional population viability analyses typically seek to identify the most sensitive life stage to environmental perturbations in order to identify management priorities, yet it is ultimately the combination of threats and their overall impact that will determine population growth rate (Beissinger and McCullough 2002). Different threats can impact different stages of an organism’s life history, and vulnerability to particular threats strongly depends on a species’ ecology (Reynolds 2003, Isaac and Cowlishaw 2004, Brooks and Kindsvater 2022). For example, in sea turtles the dominant threats posed to early life stages (e.g., nest predators, beach traffic, artificial lights) are completely different from the primary threats to adults (e.g., boat collisions, fishing bycatch; Donlan et al. 2010, Bolten et al. 2011). Similarly, in pond-breeding amphibians an increased frequency of droughts is likely to reduce recruitment success (Taylor et al. 2006, Crawford et al. 2022), whereas deforestation impacts the survival of terrestrial juveniles and adults (Todd et al. 2014). Clearly, for the effective conservation and recovery of species in the modern era, multiple novel threats experienced across a species’ life history often must be identified and evaluated.

The Eastern Hellbender (*Cryptobranchus alleganiensis,* hereafter hellbender) exemplifies the challenges associated with population assessments of long-lived, highly fecund vertebrates that simultaneously face multiple anthropogenic threats. Hellbenders are fully aquatic salamanders in the family Cryptobranchidae that live up to 30 years and produce hundreds of eggs each year after reaching sexual maturity (Petranka 1998). Historically, hellbenders inhabited streams across 15 states in the eastern United States, but they have experienced rapid local extirpations throughout their range and the reasons remain obscure (Wheeler et al. 2003, Jachowski and Hopkins 2018; USFWSg, 2018). Population declines are often associated with loss of upstream riparian forest cover and characterized by a shift in age class distribution where small populations are primarily comprised of old adults, with little evidence of recruitment of young age classes (Wheeler et al. 2003, Jachowski and Hopkins, 2018). One hypothesis for this pattern is a reduction in the number of suitable nesting sites under boulders due to high siltation caused by upstream deforestation. Alternatively, nest sites may be plentiful, but conditions in the stream may elevate rates of nest failure or larval mortality or increase the strength of density dependent interactions of larvae due to siltation of their microhabitat. Hellbenders are particularly interesting in this regard because they exhibit extended parental care of eggs and larvae (Hopkins et al. 2023), which in theory could improve the fate of nests and increase early larval survival.

However, attending males also exhibit whole-clutch filial cannibalism (Hopkins et al. 2023). Lastly, some experts noted concerning rates of adult mortality as a result of fishing and/or collection for the pet trade (Nickerson and Briggler 2007, USFWS, 2018). These hypotheses are not mutually exclusive; it is possible that some combination of these threats is generating the observed declines (Foster et al. 2009, Burgmeier et al. 2011, Unger et al. 2013, Jachowski and Hopkins 2018, USFWS 2018, Hopkins et al. 2023).

Here we develop a demographic model to understand how multiple co-occurring threats impact the population viability of a long-lived, highly fecund vertebrate with parental care. We simulate age-structured population dynamics of hellbenders using recent data on reproductive ecology collected from artificial nest boxes (Hopkins et al. 2023). Our parameterization explicitly models the fates of individual nests and includes multiple sources of density-dependent mortality – one during the egg and hatchling stages experienced within nests, and another for free-swimming larvae following emergence from nests. We explore scenarios with 1) limited nest site availability; 2) increased rates of nest failure; 3) a range of strengths of density dependence in juvenile survival rates; and 4) stochastic adult mortality. In each scenario, we quantify extinction probabilities and compare our predictions to recent evidence for changes in age class structure within declining populations (Jachowski and Hopkins 2018). We interpret our findings in the broader context of detecting declines in long-lived vertebrates with parental care and discuss how our approach can be generalized to evaluate the population viability of species facing multiple concurrent anthropogenic threats.

## METHODS

### Study Species

Hellbenders exhibit a suite of life history traits that are typical of many long-lived, highly fecund vertebrates. Individuals grow to a maximum length of approximately 570 mm (Taber 1975, Wiggs 1977), and both sexes can live for 30 years (Taber et al. 1975, Peterson et al. 1983, Makowsky et al 2010, Browne et al. 2014). Males reach sexual maturity at 4-6 years of age whereas females mature at 5-7 years of age (Figure 1c; Nickerson and Mays 1973, Taber et al. 1975, Topping and Ingersol 1981, Peterson et al. 1983). Hellbender clutch sizes generally range from 100 to 600 eggs (>90% of clutches observed in wild populations fall within this range, Hopkins et al. 2023) and eggs produced by females are positively correlated with their body size (Smith 1907, Topping and Ingersol 1981).

**Figure 1.**
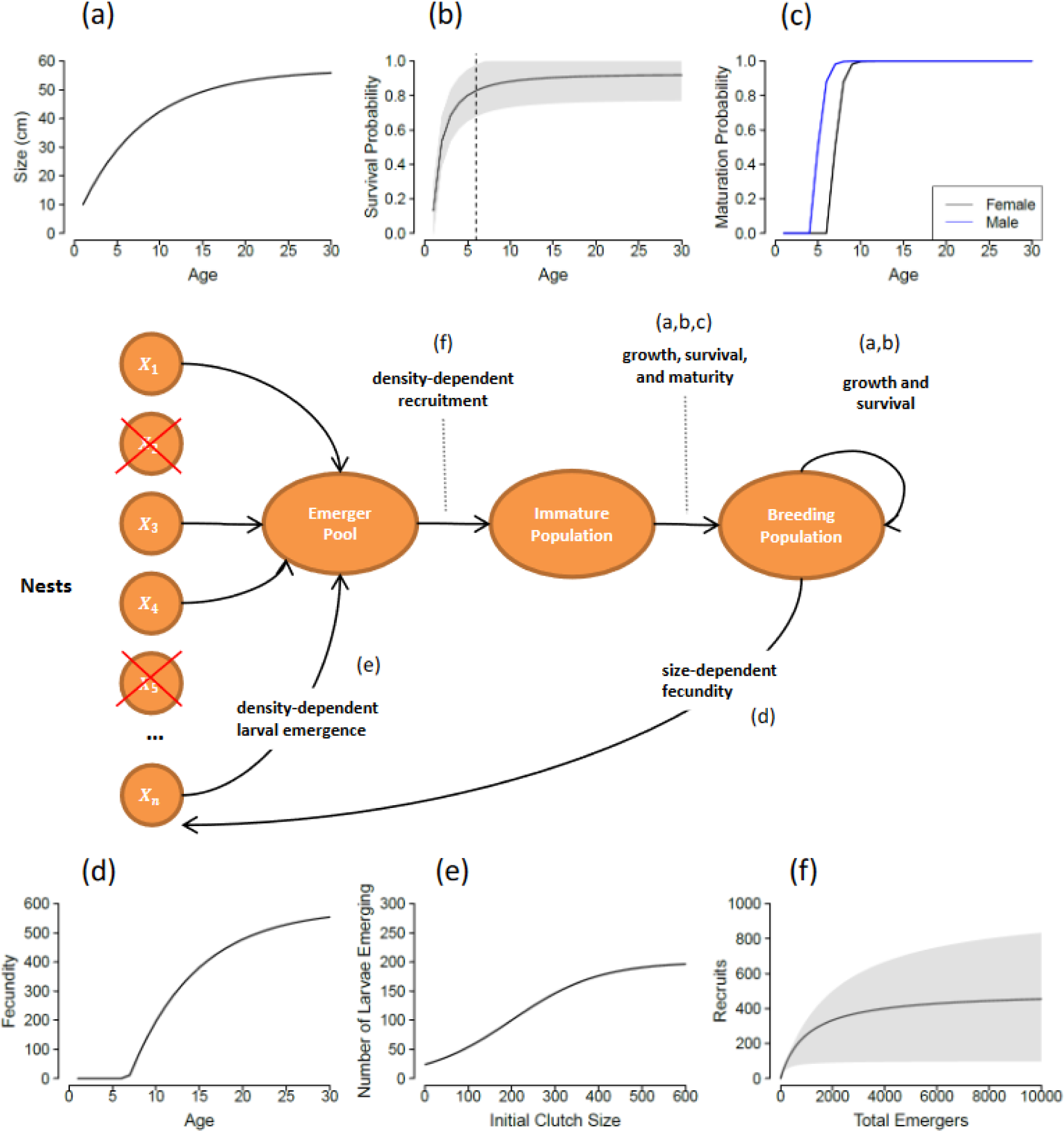
Modelled hellbender life cycle with associated vital rate functions. Nests are modelled individually, and females deposit one clutch per nest site such that the number of nest sites limits the number of clutches that can be deposited in a given year. *X*_*n*_ represents the clutch size deposited in nest site *n* in any given year, and red crosses indicate nests that fail in that year. Panels on the top row describe functions for size-dependent growth (a), survival (b), and maturation (c). Panels on the bottom row describe the relationship between clutch size and female body size (d), the relationship between initial clutch size and number of larvae emerging from nests (e), and the strength of density-dependent recruitment (f). For models that include stochasticity in survival and recruitment, the range of values drawn from is indicated by the shaded region (b, f).

Hellbenders nest under large boulders in the streams they inhabit and exhibit obligate paternal care with males guarding clutches and larvae for approximately eight months (Hopkins et al. 2023). Caring male hellbenders exhibit filial cannibalism, where they consume some or all of their clutch. Filial cannibalism is presumably a natural behavior in hellbenders, as it is relatively common in species with obligate paternal care (Manica 2002, Klug et al. 2006, Klug and Bonsall 2007). However, recent evidence from studies of hellbenders nesting in artificial shelters suggests that nest failure rates are highest at sites with low riparian forest cover, and that whole-clutch filial cannibalism is the most common reason for failure at these sites (Hopkins et al. 2023). The relative influences of nest success, nest site availability, and recruitment on population viability of hellbenders is unclear. It is likely that the longevity of adults can buffer populations against diminished reproductive returns (Chesson and Warner 1981, Chesson 1984, Taylor et al. 2006), but to what degree is unknown. To answer these questions in a modelling framework, we used specific relationships regarding growth, mortality, and size-dependent fecundity to understand the connection between demographic structure and viability.

### Growth and mortality

We used published estimates of somatic growth rates for adult hellbenders to relate total length (mm) to age (years) with a von Bertalanffy growth curve (Smith 1907, Bodinof et al. 2012, Unger and Mathis 2013). We assumed that both sexes grow at equal rates, achieve the same asymptotic body size, and experience the same size-dependent mortality rates (Figure 1b). We assume individuals face a density-dependent bottleneck in survival during the first years of life post-emergence, as young-of-the-year presumably must compete for hiding spaces and food. Although there is limited information on density-dependent processes in wild populations, siltation is often touted as the biggest threat to hellbender habitats (USFWS 2018); siltation is likely to impact populations by reducing nest sites and larval habitat, and thus we expect density-dependent survival early in life to be an important consideration even in small and declining populations. We modeled survival to the end of the juvenile period as a saturating function of density, following the Beverton-Holt recruitment function commonly used to represent recruitment of larval stages (Table 1; Kindsvater et al. 2016). We assumed an equal sex ratio at recruitment based on prior studies (Peterson et al. 1988, Humphries and Pauley 2005), although male-skewed sex ratios have been documented in some populations (Smith 1907, Hillis and Bellis 1971, Foster et al. 2009, Burgmeier et al. 2011). At age one, we assume individuals are 100 mm long (Smith 1907, Taber 1975, Peterson et al. 1983). From year one onward, we modeled hellbender survival as a function of size that increased asymptotically following the survivorship curves provided in Taber (1975), Peterson et al. (1983, 1985) and Olson (2013) (Figure 1b).

**Table 1.** Population model functions and their interpretation.

| Process | Equation | Interpretation |
| --- | --- | --- |
| **Von Bertalanffy growth function** | $L(i, a + 1) = L(i, a)e^{-k} + L_{\infty,i}(1 - e^{-k_i}) .$ | $L_{\infty,i}$ and $k_i$ determine size-at-age $L$ (for each sex $i$ at age $a$) |
| **Maturation probability function** | $p_m(i, a) = \frac{1}{1 + e^{-q(a - a_{mat,i})}}$ | Function describing the cumulative probability (ogive) individuals are mature at age $a$; $a_{mat,i}$ is the age at which 50% of individuals mature; $q$ determines the steepness of the sigmoidal curve |
| **Size-dependent fecundity** | $f(i, a) = mL(i, a) - c$ | Fecundity depends on length, which depends on age. Size-specific fecundity parameters are $m$ and $c$ |
| **Initial number of eggs per nest site, incorporating nest failure** | $X_n(t) = f(i, a) \times \text{Bernoulli}(1 - p_{fail})$ | The size of the clutch deposited in nest site $n$ at time $t$ is modelled as a function of the size of female $i$ assigned to nest site $n$. Assignments are drawn at random from the pool of mature females and nests fail with probability $p_{fail}$ |
| **Density-dependent larval emergence** | $P_n(t) = \frac{K}{1 + e^{-r(X_n(t) - K)}}$ | Where $P_n(t)$ is the number of larvae emerging from nest $n$ at time $t$, with the relationship between initial number of eggs in a nest and number of larvae emerging modelled as a logistic curve with parameters $K$ and $r$. |
| **Density-dependent recruitment** | $N_0(t + 1) = \frac{\alpha \sum P_n(t)}{1 + \beta \sum P_n(t)}$ | Beverton-Holt recruitment function |
| **Size-dependent mortality** | $M_L(i, a) = M_u L(i, a)^b$ | Mortality depends on length, which depends on age. Following Lorenzen (1996), size-specific mortality parameters are $M_u$ and $b$. This function is used to calculate the survival function in Figure 1b. |
| **Population dynamics through time** | $N(a + 1, t + 1) = \begin{cases} N_0(t) & \text{if } a = 0 \\ N(a, t)e^{-(M(a) + F(a))} & \text{if } a > 0 \end{cases}$ | $N(i, a, t)$ is the number of individuals age $a$ alive at time $t$. $N_0(t)$ is made up of equal numbers of males and females. |

### Reproduction and nest fates

Our model incorporates several key aspects of the reproductive biology of hellbenders: size-based maturation and fecundity (egg production), varying availability of nest sites, and varying nest fates. We assume that individuals mature as a probabilistic function of age, and annual egg production depends on female body size (Figure 1d). We model clutches individually to incorporate hellbender-specific nesting behavior and characterize individual nest fates. We assume that hellbender population dynamics within a stream can be represented by a core subpopulation that roughly corresponds to a 1-km reach and explore scenarios where nest sites are severely limited by siltation (nest sites < 5), and scenarios when nest sites are plentiful in that area (Jachowski and Hopkins 2018, USFWS 2018, Hopkins et al. 2023). We arbitrarily set an upper bound of 30 nest sites because exploratory analyses showed that increasing nest site availability further had minimal impact on population trends, owing to the density-dependent mortality experienced by larvae post-emergence (Appendix S1: Section S3). In our model, females are randomly assigned to a nest site. If there are more females than nest sites in a particular year/scenario, females that are not randomly selected do not breed in that year. For simplicity, we did not consider size-based competition among males or females for access to nest sites, or the case where multiple females laid clutches at the same nest site. Each female that is assigned to a nest site deposits a clutch of eggs, and the size of the clutch is determined by the body size of the female (Figure 1). Because hellbenders exhibit obligate paternal care, all clutches laid are randomly assigned to a male; if there are fewer males than clutches in a given year/scenario, clutches that are not assigned to a male are eliminated. We assume that paternal care behavior and the probability of cannibalism of eggs does not vary among males of different sizes or ages, based on recent work (Hopkins et al. 2023).

### Larval emergence

After a clutch is laid in early autumn, it takes eight months for surviving hellbender larvae to emerge. The caring male occupies the nest site over winter; based on empirical observations, male presence appears to be essential for nest success. Observations of wild populations have documented nest failure due to predation, abandonment, environmental factors such as flooding, or whole-clutch cannibalism by parental males (Hopkins et al. 2023). The proportion of nests failing varies considerably across populations (Hopkins et al. 2023); therefore, we considered a range of nest failure rates in our model scenarios. In each scenario, a random sample of nests are assumed to fail and removed from the population before emerging larvae are pooled and subjected to density-dependent recruitment (Figure 1f). The proportion of the nests that fail varies with the background rate of nest failure *pfail* (Table 2); the number of larvae that die with the loss of a nest varies depending on the size of the female that laid a clutch at that nest site. Recent empirical evidence suggests the number of larvae emerging from nests is positively correlated with initial clutch size in small clutches but shows no relationship in large clutches (Hopkins et al. 2023). We therefore modeled the number of emerging larvae per nest as an asymptotic function of the number of eggs laid in a successful nest (Figure 1e).

**Table 2.**
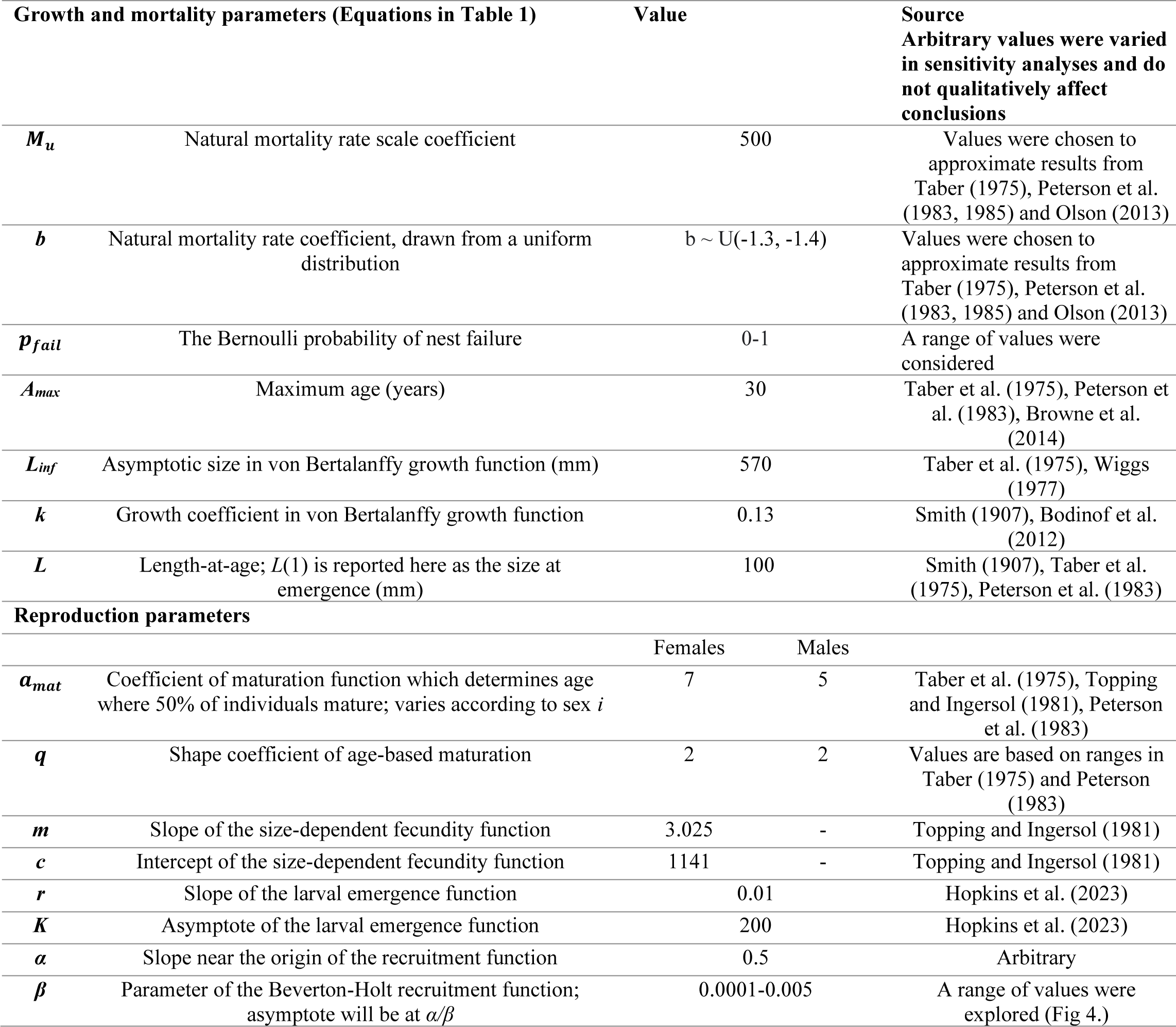
Modelled life-history parameters and their sources.

### Modelling Framework

We developed a size- and age-structured population model that includes the key aspects of the hellbender life cycle described above. Our approach is based on a model of population demography and growth of fishes with obligate paternal care in Kindsvater et al. (2020), which itself is adapted from the models commonly used in stock assessments of fished populations (Mangel 2006). However, we significantly expand the scope of this previous model to include biologically realistic processes that could affect hellbender population dynamics, such as variation in availability of appropriate nest sites, two sources of density-dependent mortality in early life stages, loss of adults from the breeding population, and environmental stochasticity (Figure 1). Our approach is distinct from previous assessments of hellbender population dynamics, which assumed a fixed proportion of larvae grew into juveniles, resulting in exponential population growth (Unger et al 2013). We started from an arbitrary population size (100 adults and 1000 juveniles) and allowed the population to equilibrate and reach a stable age distribution, representing a plausible baseline for its historic age structure and size. After emerging from the nest, juveniles spend the rest of the year at large before recruiting to the age-structured population, a process represented with a density-dependent recruitment function (Table 1). We projected population dynamics under different scenarios representing the individual and interactive effects of nest site availability, nest failure predominantly due to filial cannibalism, and variability in adult mortality, in order to understand their relative contributions to population demography and viability into the future. All functions and parameters representing each process are in Tables 1 and 2. All models were programmed in R (R Core Team, 2023) and the code needed to reproduce the analysis is archived on Zenodo (Brooks et al. 2023).

### Deterministic simulations

We first simulated populations with rates of nest failure ranging from 0 – 80% and allowed each population to reach equilibrium. Because nest failure in artificial cavities can reach 100% in degraded sites (Hopkins et al. 2023), we then increased nest failure rate for each scenario to evaluate the demographic consequences of chronic recruitment failure. In order to isolate the effect of nest failure rates on population growth, we assumed that nest sites were not limiting (n = 30), and adult survival was constant in these initial scenarios. Specifically, the *b* parameter in the size-dependent mortality function was fixed at −1.38 (Table 1). Further, we treated juvenile survival and recruitment rates as fixed parameters in order to investigate deterministic population dynamics and demography. Specifically, α and β parameters of the Beverton-Holt recruitment function were set to 0.5 and 0.001, respectively (Table 1). Essentially, this approach assumes that spatial and temporal variation average out over the total population, such that mean parameter values provide good approximations of the fate of a given cohort. To explore the contributions of density dependence at different life stages, we compared simulations with and without density-dependent larval emergence, and with and without Beverton-Holt recruitment.

### Stochastic simulations

Environmental and demographic stochasticity are not captured in our deterministic simulations, yet are an important determinant of extinction risk, particularly for small populations (Lande 1993). In order to incorporate the effects of interannual variation in environmental conditions (e.g., stream flows / temperatures) and the role of chance in the fate of small populations, we next implemented stochastic survival functions at multiple life stages to estimate average time to extinction of populations under different scenarios (Figure 1). Survival was partitioned between mature and immature animals to separately model fluctuations in recruitment success and sporadic harvest of breeding adults. For each year of the simulations, survival probabilities were randomly drawn from a uniform distribution of plausible values. We compared models where only the survival of immature individuals post-emergence varied across years with models where survival of all individuals varied (Table 2, Figure 1b). For both survival scenarios we explored the full range of nest failure rates (0-100%) and variation in the number of available nest sites (0-30) to understand the interacting contributions of these processes. To account for uncertainty in the fate of larvae immediately following emergence from nests, we explored scenarios with varying strengths of density-dependent recruitment by modifying the beta parameter (Table 2, Figure 1f). We ran 1000 simulations of each scenario and tallied the number of simulations that dropped below an arbitrary threshold (<5 adult females) over a 200- year projection. Dividing these tallies by 1000 gave the probability of quasi-extinction for each combination of nest site availability, nest failure rate, and adult mortality.

## RESULTS

### Deterministic simulations

We found that hellbender populations are predicted to persist under a broad range of nest fates (0-80% failure) when nest site availability and adult survival are both high (Figure 2), although the rate of nest failure determines the equilibrium abundance. Nest failure rates of 90 and 99% however, result in population declines (Figure 2). We examined the population age structure over several time intervals for both the 90% and 99% nest failure scenarios (Figure 3); in both cases the long-lived adults persisted for many years after the introduction of chronic recruitment failure, but populations dwindled toward a geriatric age-structure over time (Figure 3). Unsurprisingly, when 99% of nests failed, it was catastrophic in all scenarios; all populations went extinct within 30 years. Although similar declines in population abundance occurred at 90% nest failure, populations persisted for over a century before complete collapse.

**Figure 2.**
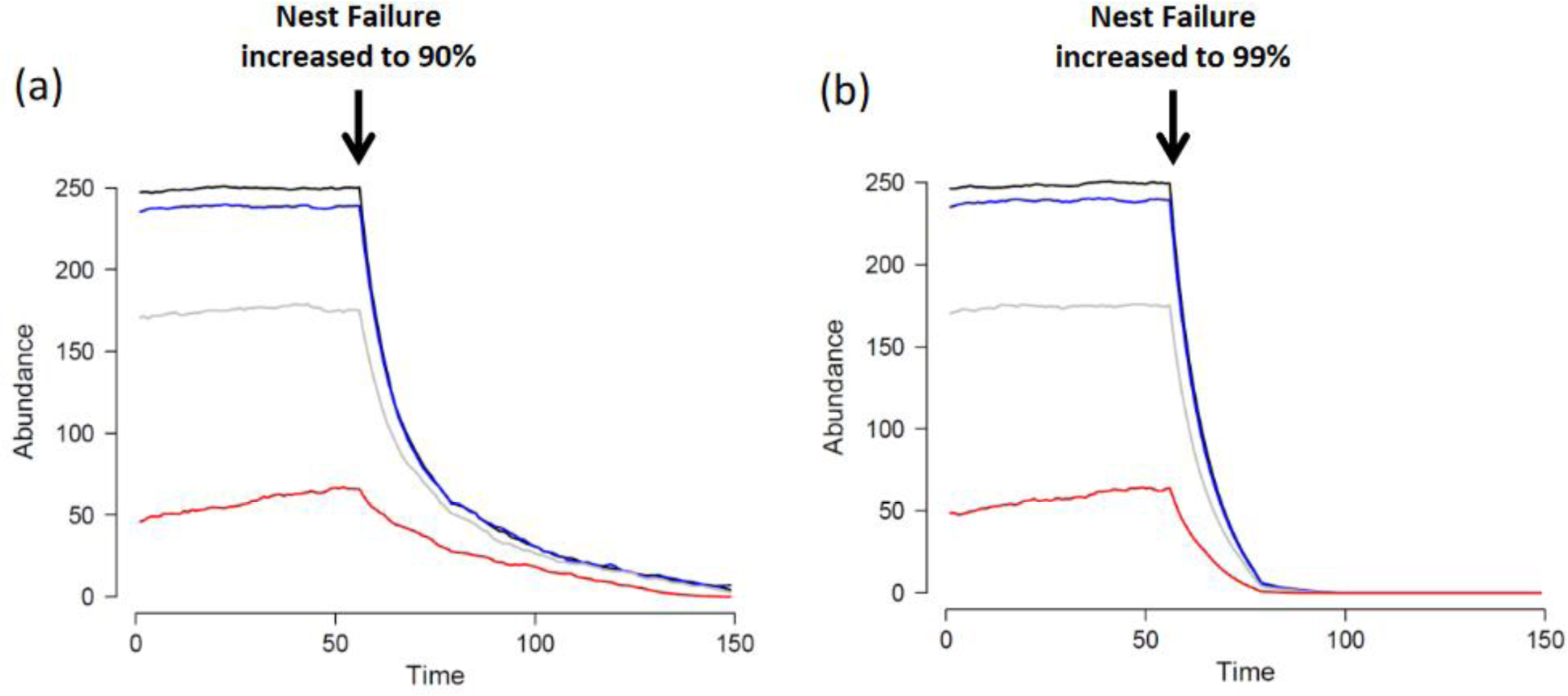
Projected abundance of adult hellbenders for various levels of nest failure (black line = 0% nest failure, blue line = 10% nest failure, grey line = 50% nest failure, red line = 80% nest failure), without nest site limitation. At 50 years, nest failure is increased to a) 90% and b) 99% for all scenarios to illustrate how populations respond to a shift in environmental conditions (e.g., deforestation of riparian buffers upstream of a population).

**Figure 3.**
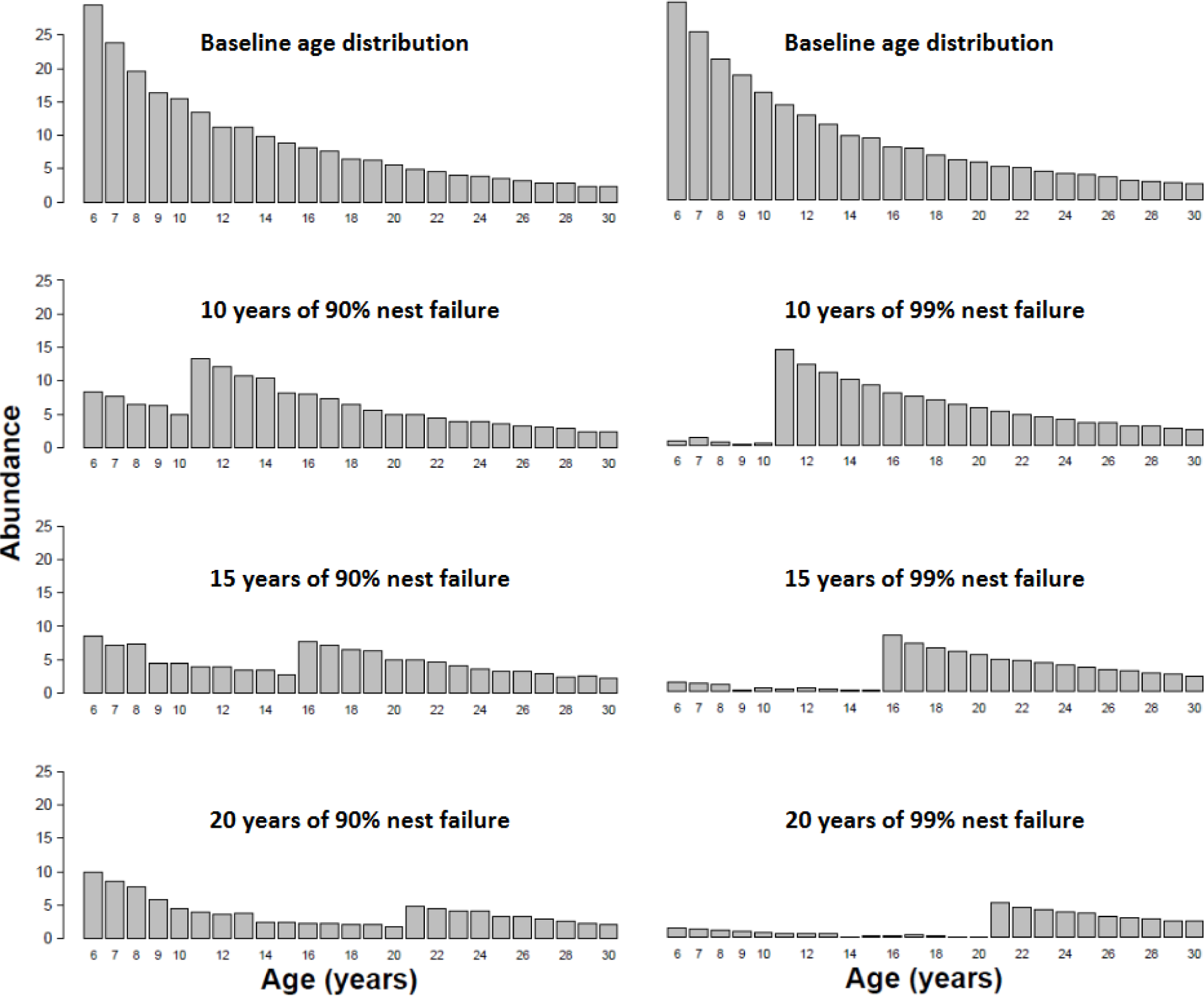
The change in adult age distribution after a sharp decline in nest success. Top row represents the stable age distribution at 50% nest failure (grey line in figure 2). The subsequent panels illustrate the age distribution 10, 15, and 20 years after a shift to 90% (left column) and 99% (right column) nest failure. Immature individuals were excluded from the figure to make it easier to visualize declines of breeding adults.

Removing the density-dependent terms in the model did not change our general conclusions (Appendix S1: Section S1, Appendix S1: Section S2). Even without density-dependent larval emergence and survival, a tipping point still exists (at ∼90% nest failure) where populations are no longer self-sustaining. Further the speed of the decline when nest failure is increased beyond this threshold is comparable in simulations with and without density dependence. However, when density-dependence is not accounted for, models yield population size estimates that are unrealistic (up to 2500 adults in a 1km stretch of stream with only 30 nests). These unrealistic estimates reinforce our conclusion that density-dependent processes – such as competition for structural habitat shelter in silted streams – are likely limiting juvenile survival in hellbender populations. Further, this finding highlights the importance of including density-dependent processes more generally; failure to account for density dependence can result in the tendency to overstate declines relative to historic abundances and generate overly optimistic forecasts regarding the rate of population recovery.

### Stochastic simulations

Nest site availability, rates of nest failure, and the strength of density-dependent recruitment (juvenile survival) all exhibited a threshold beyond which extinction was certain (Figure 4). In most instances, stochastic adult survival increased extinction probability relative to the equivalent deterministic scenario, and ultimately reduced the range of conditions under which populations were able to persist. Unsurprisingly, extinction probabilities increased with increasing rates of nest failure in a similar fashion to the deterministic simulations but exhibited a lower threshold beyond which extinction became inevitable (∼80% vs. 90% failure; Figure 2, 4). However, this robustness to extinction only held when nest availability was high; with fewer nests overall, extinction was inevitable at lower nest failure rates (∼40%; Figure 4, Appendix S1: Section S3). In other words, when nest failure rates were high, limited availability of nest sites (<10 nest sites available) was an important contributor to extinction risk (Figure 4a, b); however, the threat posed by limited nest site availability was negligible if nest success was high. Extinction risk also increased as the strength of density-dependent recruitment increased, with a tipping point occurring at moderate levels of density dependence (Figure 4c, d). Similarly, our analysis shows that the effect of sporadic adult mortality was most severe when most nests failed, and substantially increased the probability of extinction within 200 years (Figure 4a, b). Sensitivity analysis confirmed that variable survival exerted a greater impact on extinction risk than the specific form of the survival function (Appendix S1: Section S4). Increased nest site availability mitigated extinction risk only to a minor extent, suggesting the longevity of adults is more important for offsetting heavy juvenile losses in such scenarios. In summary, high rates of nest failure and/or juvenile mortality can lead to population collapse in our models. The situation is most severe when adult mortality is high or, to a lesser extent, when nest sites are limited, because these act to critically undermine the population’s ability to buffer against natural variation in recruitment.

**Figure 4.**
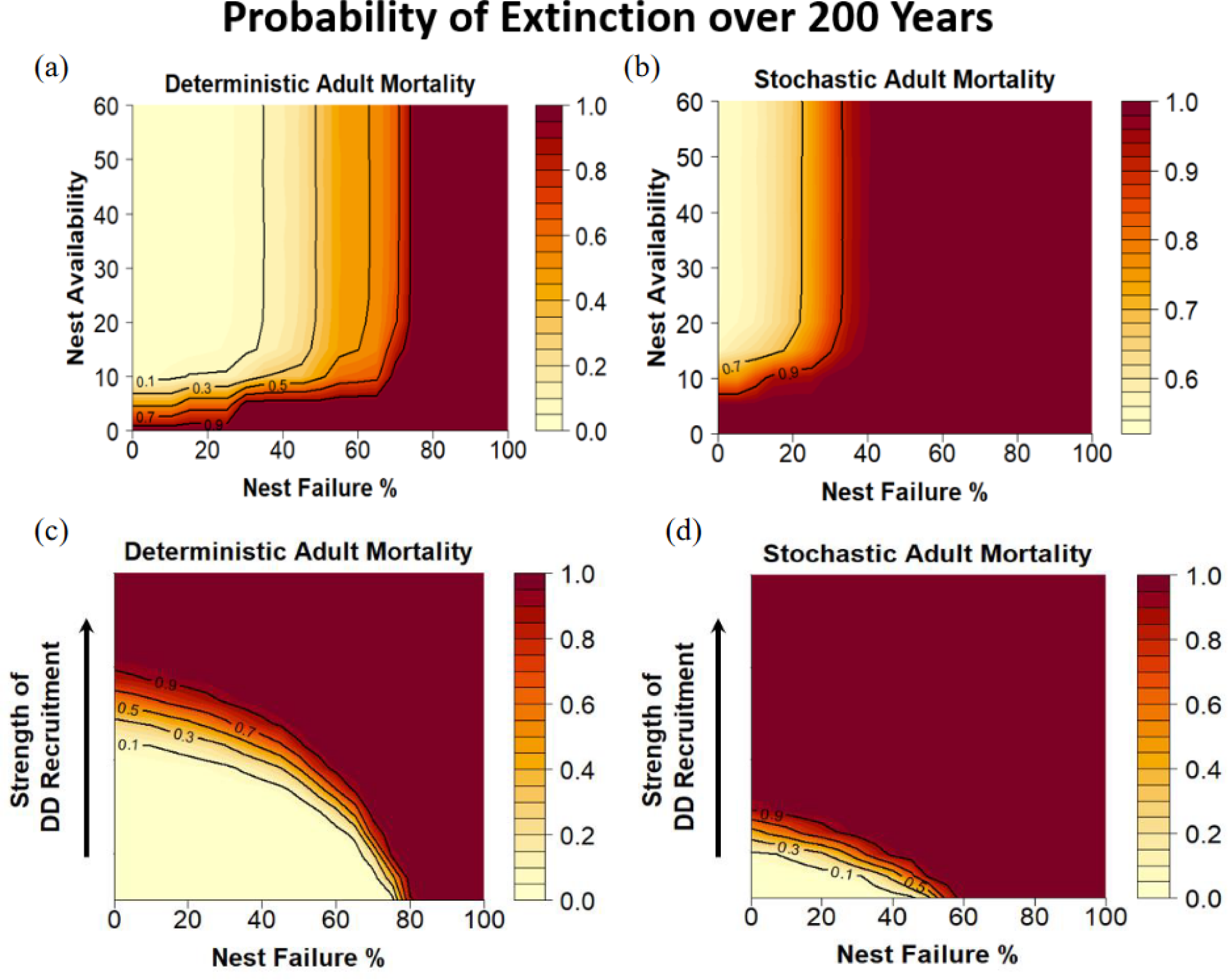
Extinction probabilities as a function of nest site availability, rate of nest failure, and strength of density-dependent recruitment. Extinction probabilities were obtained from 10,000 simulations projected over 200 years for each combination of factors. Darker shades indicate higher probabilities of extinction. Every scenario was simulated twice under deterministic (a, c) and stochastic (b, d) adult mortality. For scenarios exploring a range of nest site availabilities (a, b), uncertainty in the strength of density-dependent larval recruitment from year to year was captured by averaging extinction probabilities across scenarios, representing stochastic recruitment. For scenarios that explored a range of density-dependent recruitment levels (c, d), nest site availability was held constant at 30 nests.

## DISCUSSION

Here we present a demographic model that incorporates key aspects of life history, particularly reproductive ecology and recruitment, to understand the population dynamics of a long-lived, highly fecund vertebrate. Our framework illustrates how multiple co-occurring threats impacting different stages of an organism’s life cycle can undermine population viability and shows how the life history of hellbenders makes them particularly susceptible to extinction debt. We demonstrated that the high rates of nest failure that have been observed in the field (Hopkins et al. 2023) are sufficient to drive hellbender populations towards a geriatric age distribution and eventually to localized extinction, but that this process takes decades due to the species’ longevity. As a result, population declines can go unnoticed for years because adults persist on the landscape. As has been shown in other long-lived taxa (Hutchings et al. 2012, Spencer et al. 2017), we also demonstrated that moderate reductions in the survival rates of mature hellbenders can threaten population persistence. In contrast, limited nest site availability alone had comparatively little influence on population dynamics due to density-dependent compensation in the juvenile stages, but limited nest sites can exacerbate the effects of excessive adult mortality and high rates of nest failure, quickly jeopardizing population viability.

Environmental stochasticity has led to the evolution of life histories that can overcome times of low productivity (Shaffer 1981, 1987, Stearns 1992, Lande 1993); long-lived, highly fecund species are relatively robust to annual variability in recruitment, and it is only typically chronic reproductive failure that will result in extinction debt and population declines (Hutchings et al. 2012). In contrast however, long-lived, highly fecund species are sensitive to annual variability in adult mortality (Spencer et al. 2017). Population growth can be sensitive to changes in adult survival rates in long-lived species, such that any novel sources of adult mortality can quickly threaten the long-term persistence of populations (Congdon et al., 1994; Taylor & Scott 1997; Biek et al. 2002; Hels & Nachman 2002; Vonesh & De la Cruz 2002, Schmidt et al. 2005, Kindsvater et al. 2016). For long lifespans to evolve, adult survival must be largely predictable (Stearns 1992), and as such it is likely that survival rates of adult hellbenders remained relatively constant for much of their evolutionary history. In the contemporary era however, novel sources of anthropogenic mortality emerged that can undermine the breeding population’s ability to replace itself. Thus, quantifying the effect of stochasticity, in terms of which life stages it influences and its impact on population growth, is essential to understand its contribution to extinction risk. Stochastic forces disproportionately threaten small populations (Lande 1993), highlighting its importance in conservation fields. As demonstrated in our simulations, annual variability in adult survival rates acts to substantially increase extinction risk. Here we show that failing to account for stochastic forces results in overestimating persistence probabilities of hellbenders (Figure 4), and likely this represents a common feature of population assessments for long-lived species with slow recovery times.

Long-lived species with high adult survival have evolved to persist with relatively low recruitment, where most juveniles do not survive to adulthood (Chesson and Warner 1981, Warner and Chesson 1985, Childs et al. 2010). As a result, the longevity of breeding adults can offset high juvenile mortality and enable population persistence, but our models show this may not be enough to prevent extinction in modified landscapes. Firstly, even though hellbenders evolved to tolerate high rates of egg and larval mortality, there are limits above which populations will decline, especially when coupled with other threats operating on different life stages. Second, the combined effects of limited nest sites and elevated rates of nest failure create a scenario of extinction debt, in which local extirpation of hellbender populations becomes likely even when adults persist on the landscape. Any one of these factors alone can to some extent be compensated for by longevity coupled with multiple bouts of reproduction but taken together they undermine a population’s ability to persist in landscapes impacted by humans. Unfortunately, knowledge concerning adult mortality is often lacking for long-lived species because the long-term studies required to obtain accurate estimates of survival rates are rare (Congdon et al. 1994). Demographic models can explore a range of scenarios in the face of such uncertainty but could be greatly improved with empirical data.

Assessments of population status require a comprehensive understanding of the focal species’ natural history. Future studies should seek to obtain such missing pieces of natural history information that may strongly influence the dynamics of declining populations. Indeed, parameterizing our hellbender models has only become possible with the recent acquisition of data on key aspects of their reproductive ecology from nest box studies (Hopkins et al. 2023). These new insights have shed considerable light on the dynamics of early life stages, the factors that govern nest outcomes, and potential causes for recent declines. There are, however, several additional aspects of breeding behavior that we have omitted due to lack of detailed evidence: 1) multiple females depositing clutches at the same nest site (which may be particularly important when nest sites are limiting), 2) competition for mates and/or nest sites (which may be size-based and reflect variation in the quality of nest sites), and 3) spatially explicit breeding dynamics. Similarly, the factors regulating larval and juvenile survival post-emergence are largely unknown for many species, including hellbenders (Chan et al. 2018). As illustrated by our models however, population dynamics of long-lived, highly fecund species are influenced strongly by our assumptions regarding density-dependent regulation. The strength of density-dependent survival in early life stages will determine the relative contribution of high fecundity to population resilience, and thus control the tipping points that lead to declines (Reynolds et al. 2005, de Silva et al. 2019). For hellbenders, increased siltation has been proposed as a mechanism that reduces the availability of habitat for newly emerged larvae (USFWS 2018), and thereby strengthens density-dependent mortality of early life stages. This idea, however, has yet to be formally tested. In fact, improved techniques for detecting hellbender larvae would enable fundamental studies of larval ecology, which remains among the most important knowledge gaps for the species (Bodinof et al. 2012, Unger and Mathis 2013). Studies specifically designed to quantify density dependence across life stages, and how they are affected by environmental change, would greatly advance our fundamental understanding of population regulation in hellbenders and other species of conservation concern.

Demographic models such as the one presented here can help to alleviate some of the challenges associated with population assessments of long-lived, highly fecund species (Beissinger and McCullough 2002). By explicitly incorporating multiple stage-specific impacts into modelling frameworks, it is possible to gain insight into the contributions of different anthropogenic threats to the extinction risk of threatened and endangered taxa. This information can then be used to diagnose cryptic declines, identify instances of extinction debt, and outline the management interventions necessary for population recovery. Despite a long history of practitioners advocating for conservation that considers all life-history stages (Congdon et al. 1994, Wake and Vredenburg 2008, Mace et al. 2018), there are still comparatively few studies that seek to quantify the combined impact of multiple, concurrent threats on the fate of wild populations. As we face unprecedented rates of environmental change, such a holistic approach will be necessary to preserve long-lived species that are disappearing throughout their range

## ACKNOWLEDGEMENTS

We thank the fantastic team of technicians and students over the last 15 years that assisted in collecting the empirical data that made parameterization of our models possible. We would also like to thank J.D. Kleopfer and M. Pinder for supporting our work. We would like to thank S. Button, N. Pacoureau, and K. Theberge for comments on an earlier draft that greatly improved the manuscript. This study was made possible with funding from the Virginia Department of Wildlife Resources (EP2443089), The National Science Foundation (IOS-1755055), the U.S. Forest Service, the Department of Fish and Wildlife Conservation, the Fralin Life Sciences Institute, the Global Change Center at Virginia Tech, and the National Fish and Wildlife Foundation.

## AUTHOR CONTRIBUTIONS

All authors conceived the ideas and designed methodology; GCB and HKK performed the analysis; GCB led the writing of the manuscript. All authors contributed critically to the drafts and gave final approval for publication.

