## Supplemental File 1 for "Concurrent threats and extinction risk in a long-lived, highly fecund vertebrate with parental care"

**[1]Department of Fish and Wildlife Conservation, Virginia Tech, 310 West Campus Drive, Blacksburg, Virginia, USA**

### Concurrent threats and extinction risk in a long-lived, highly fecund vertebrate with parental care

**Ecological Applications**

### Section S1: Sensitivity analysis of density-dependent recruitment

Here we provide the results from the deterministic analysis of the main text in the absence of density-dependent recruitment. The equilibrium sizes of simulated populations are approximately four times larger when density-dependence is removed (Figure S1), but when nest failure is raised to 99% simulated populations show the same shift in demographic age structure and similar times to extinction (Figure S1, S2).

Table S1. Recruitment parameters used in supplemental analyses 1. The parameters only differ from those in the main text with respect to the omission of the Beverton-Holt density dependent recruitment function.

| Reproduction parameters | | Value | |
|---|---|---|---|
| | | Females | Males |
| $a_{mat}$ | Coefficient of maturation function which determines age where 50% of individuals mature; varies according to sex | 7 | 5 |
| $q$ | Shape coefficient of age-based maturation | 2 | 2 |
| $m$ | Slope of the size-dependent fecundity function | 3.025 | - |
| $c$ | Intercept of the size-dependent fecundity function | 1141 | - |
| $r$ | Slope of the larval emergence function | 0.01 | |
| $K$ | Asymptote of the larval emergence function | 200 | |
| $\alpha$ | Slope near the origin of the recruitment function | NA | |
| $\beta$ | Parameter of the Beverton-Holt recruitment function; asymptote will be at $\alpha/\beta$ | NA | |

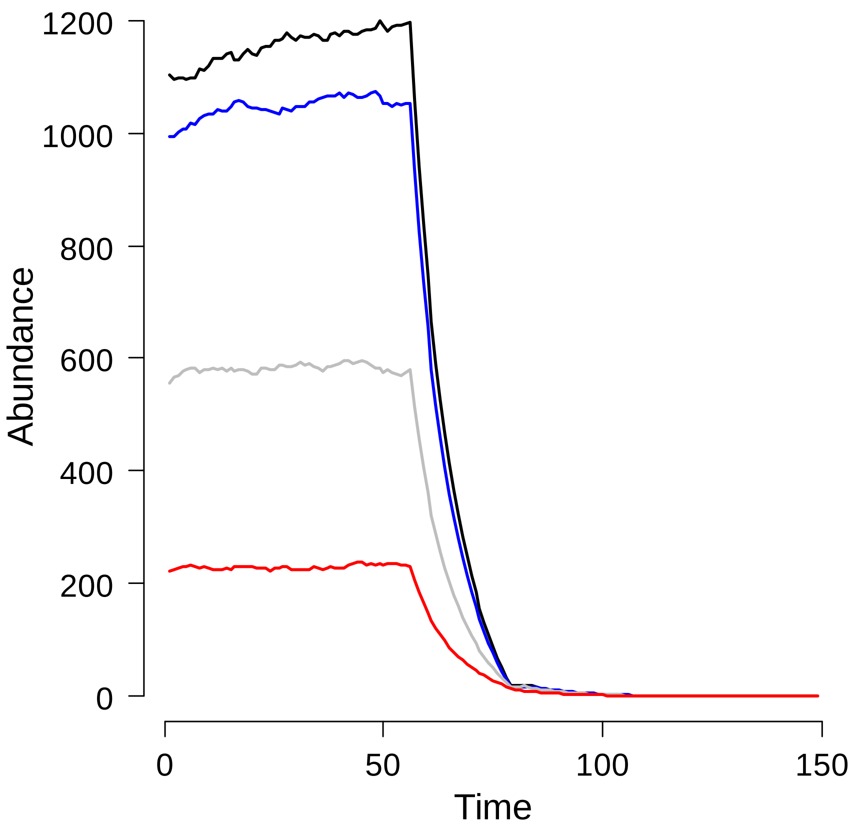


Figure S1. Projected abundance of adult hellbenders for various levels of nest failure (black line = 0% nest failure, blue line = 10% nest failure, grey line = 50% nest failure, red line = 80% nest failure), without nest site limitation or density-dependent larval emergence. At 50 years, nest failure is increased to a) 90% and b) 99% for all scenarios to illustrate how populations respond to a shift in environmental conditions (e.g., deforestation of riparian buffers upstream of a population).

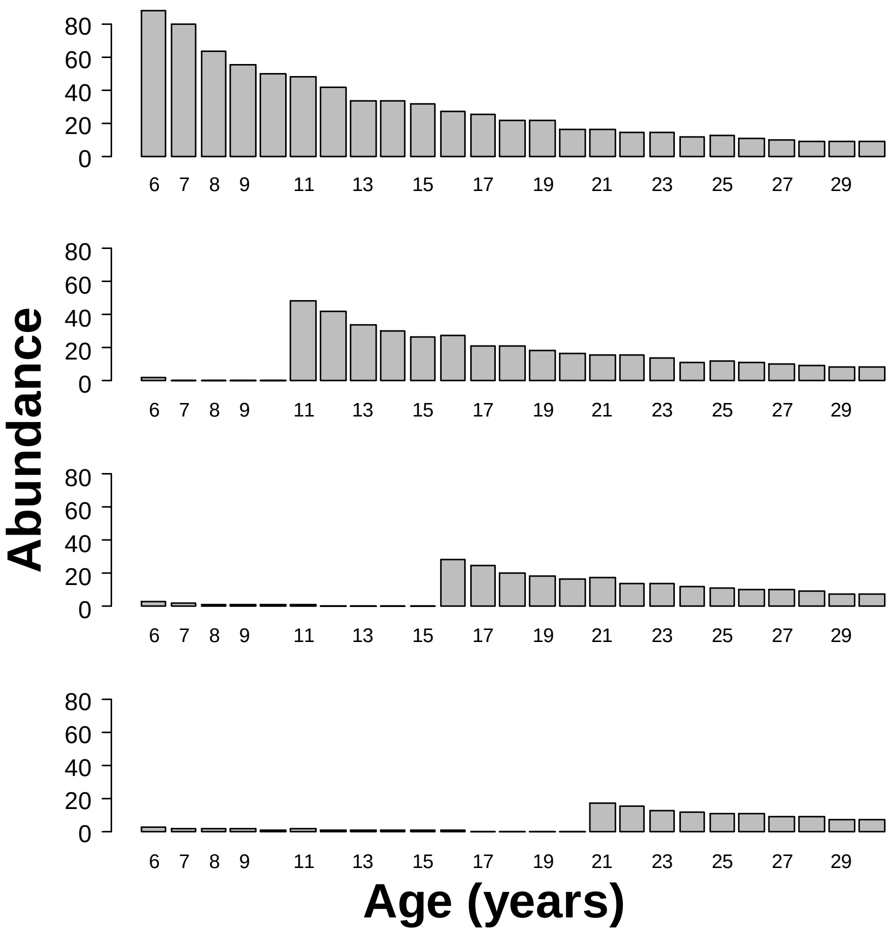


Figure S2. The change in adult age distribution after a sharp decline in nest success. Top row represents the stable age distribution at 50% nest failure (grey line in figure 2). The subsequent panels illustrate the age distribution 10, 15, and 20 years after a shift to 90% (left column) and 99% (right column) nest failure. Immature individuals were excluded from the figure to make it easier to visualize declines of breeding adults.

### Section S2: Sensitivity analysis of density-dependent larval emergence and recruitment

Here we provide the results from the deterministic analysis of the main text in the absence of density-dependent larval emergence and recruitment. The equilibrium sizes of simulated populations are approximately ten times larger when both sources of density-dependence are removed (Figure S3), but when nest failure is raised to 99% simulated populations show the same shift in demographic age structure and similar times to extinction (Figure S3, S4).

Table S2. Recruitment parameters used in supplemental analyses 2. The parameters only differ from those in the main text with respect to the omission of density-dependent larval emergence and the Beverton-Holt recruitment function.

| **Reproduction parameters** | | **Value** | |
|---|---|---|---|
| | | Females | Males |
| $a_{mat}$ | Coefficient of maturation function which determines age where 50% of individuals mature; varies according to sex | 7 | 5 |
| $q$ | Shape coefficient of age-based maturation | 2 | 2 |
| $m$ | Slope of the size-dependent fecundity function | 3.025 | - |
| $c$ | Intercept of the size-dependent fecundity function | 1141 | - |
| $r$ | Slope of the larval emergence function | NA | |
| $K$ | Asymptote of the larval emergence function | NA | |
| $\alpha$ | Slope near the origin of the recruitment function | NA | |
| $\beta$ | Parameter of the Beverton-Holt recruitment function; asymptote will be at $\alpha/\beta$ | NA | |

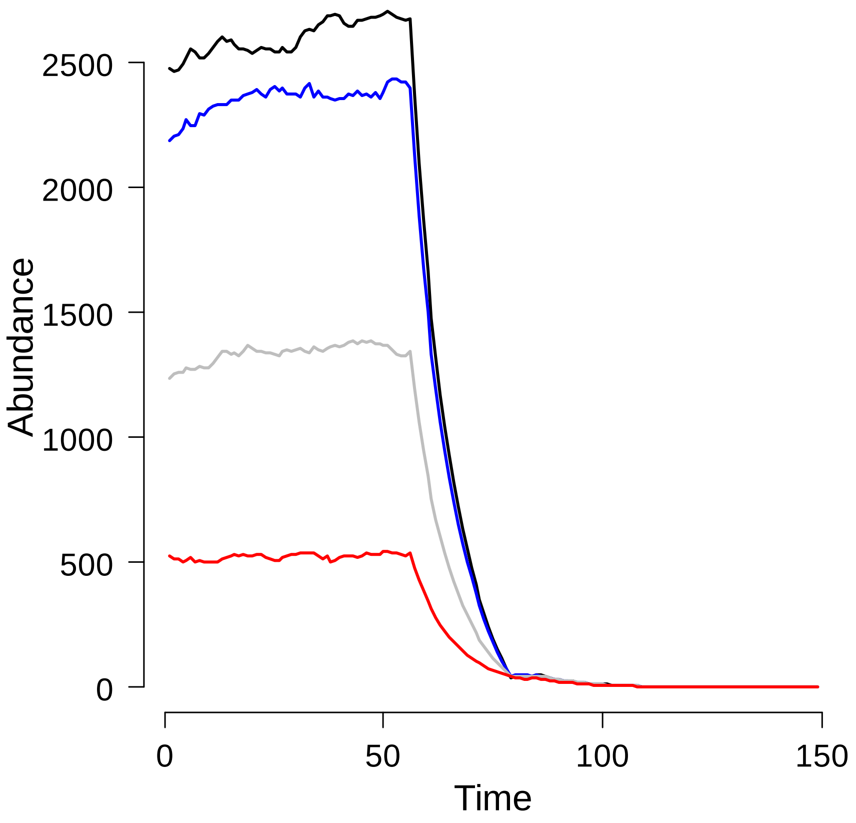


Figure S3. Projected abundance of adult hellbenders for various levels of nest failure (black line = 0% nest failure, blue line = 10% nest failure, grey line = 50% nest failure, red line = 80% nest failure), without nest site limitation, density-dependent larval emergence, or density-dependent recruitment. At 50 years, nest failure is increased to a) 90% and b) 99% for all scenarios to illustrate how populations respond to a shift in environmental conditions (e.g., deforestation of riparian buffers upstream of a population).

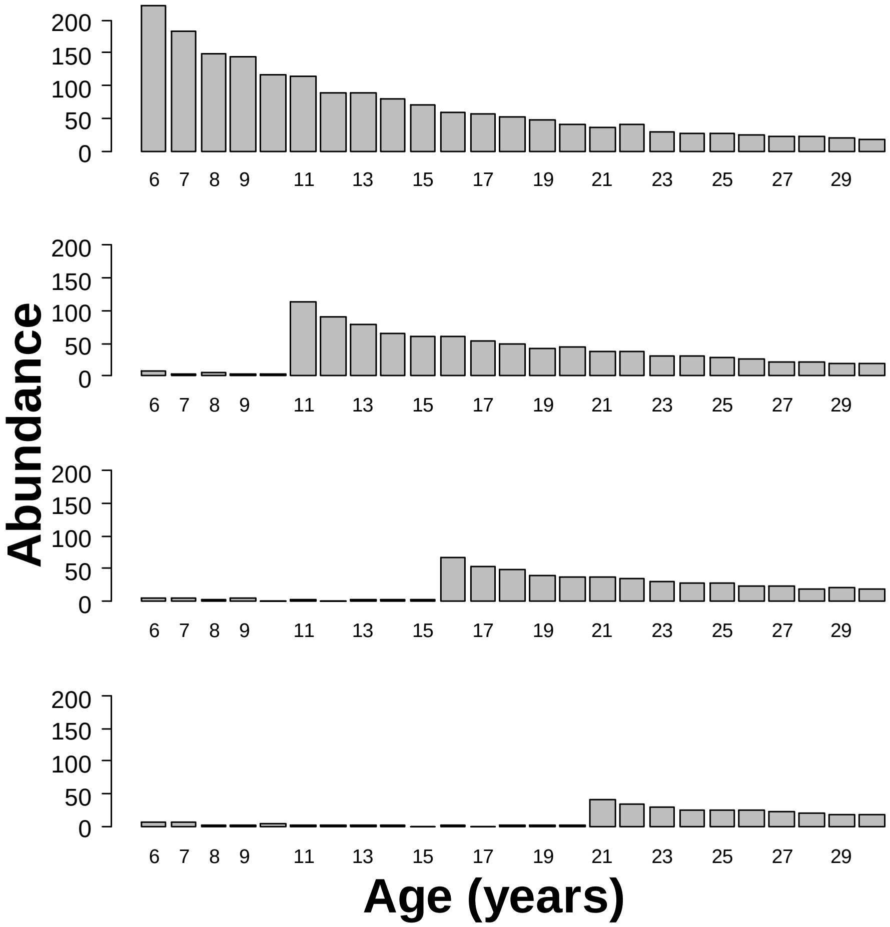


Figure S4. The change in adult age distribution after a sharp decline in nest success. Top row represents the stable age distribution at 50% nest failure (grey line in figure 2). The subsequent panels illustrate the age distribution 10, 15, and 20 years after a shift to 90% (left column) and 99% (right column) nest failure. Immature individuals were excluded from the figure to make it easier to visualize declines of breeding adults.

### Section S3: Sensitivity analysis of nest availability

Here we provide the results from the stochastic analysis of the main text for a broader range of nest availability. Owing to the presence of density-dependent larval survival, increasing the number of nests beyond a certain threshold has no impact on extinction probabilities.

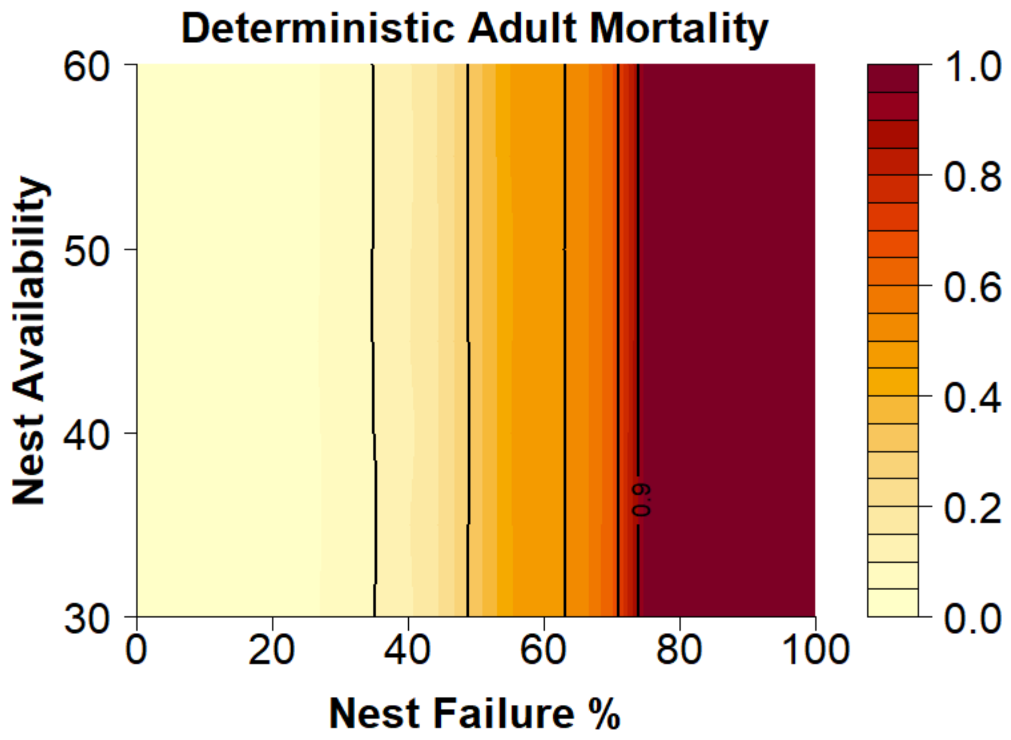


Figure S5. Extinction probabilities as a function of nest site availability, rate of nest failure, and strength of density-dependent recruitment. Extinction probabilities were obtained from 10,000 simulations projected over 200 years for each combination of factors. Darker shades indicate higher probabilities of extinction. Parameter values were identical to those used to produce Figure 4A in the main text, except nest availability ranged from 30 to 60.

### Section S4: Sensitivity analysis of survival function

Here we provide the results from the stochastic analysis of the main text but hold survival constant with respect to body size. This alternative parameterization highlights that the specific shape of the survival function has a smaller impact on extinction probabilities compared to the inclusion of variable survival across years.

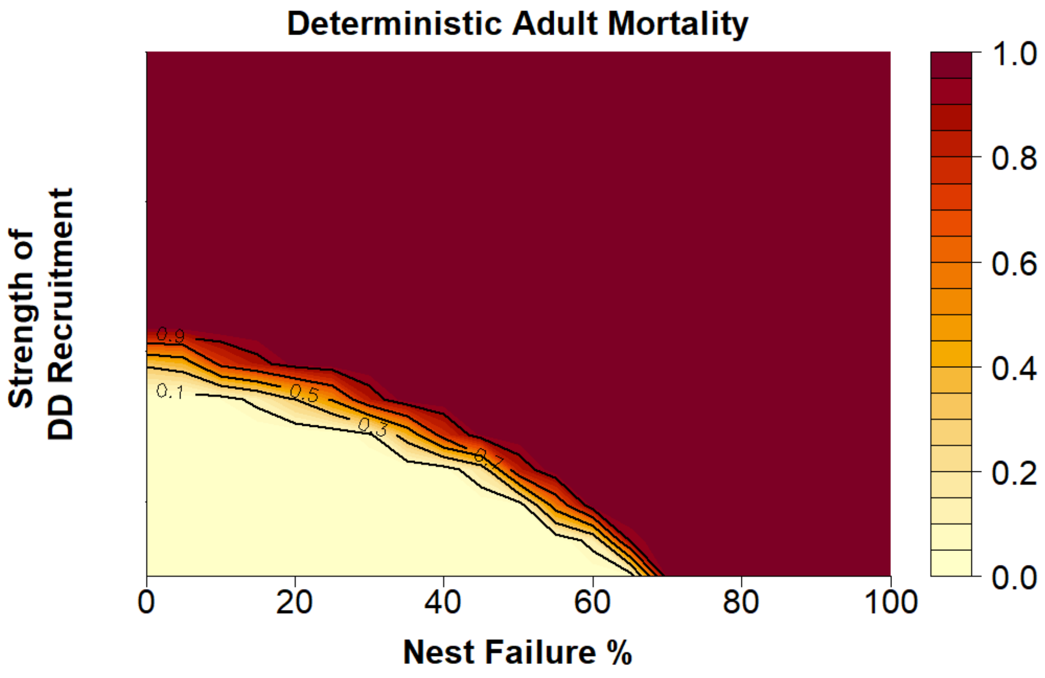


Figure S6. Extinction probabilities as a function of nest site availability, rate of nest failure, and strength of density-dependent recruitment. Extinction probabilities were obtained from 10,000 simulations projected over 200 years for each combination of factors. Darker shades indicate higher probabilities of extinction. Parameter values were identical to those used to produce Figure 4C in the main text, except survival was held at constant values (juvenile = 0.6, adults = 0.9) rather than varying as a function of body size.
